# Columba: Fast Approximate Pattern Matching with Optimized Search Schemes

**DOI:** 10.1101/2025.03.26.645543

**Authors:** Luca Renders, Lore Depuydt, Travis Gagie, Jan Fostier

**Author notes:** The authors wish it to be known that, in their opinion, the first two authors should be regarded as joint first authors.

## Abstract

Aligning sequencing reads to reference genomes is a fundamental task in bioinformatics. Aligners can be classified as lossy or lossless: lossy aligners prioritize speed by reporting only one or a few high-scoring alignments, whereas lossless aligners output all optimal alignments, ensuring completeness and sensitivity. This paper introduces Columba, a high-performance lossless aligner tailored for Illumina sequencing data. Columba processes single or paired-end reads in FASTQ format and outputs alignments in SAM format. By utilizing advanced search schemes and bit-parallel alignment techniques, Columba achieves exceptional speed. Columba is available in two variants. The first is based on the bidirectional FM-index. The second, Columba RLC, employs run-length compression using a bidirectional move structure, significantly reducing memory usage for large, repetitive datasets like pan-genomes. Through extensive benchmarking, Columba outperforms existing lossless aligners in speed, particularly at higher error rates. Tests on the human genome and bacterial and human pan-genome datasets demonstrate Columba’s robustness and efficiency. We integrated Columba into the OptiType HLA genotyping pipeline, where it substantially reduced computational time while maintaining accuracy. These results position Columba as a versatile, state-of-the-art tool for high-sensitivity genomic analyses.

## Introduction

Sequence alignment is a fundamental task in bioinformatics, with applications in variant calling, genome assembly, transcriptome analysis, functional genomics, and more. Alignment tools can be broadly categorized as lossy or lossless (see Table 1).

**Table 1.**
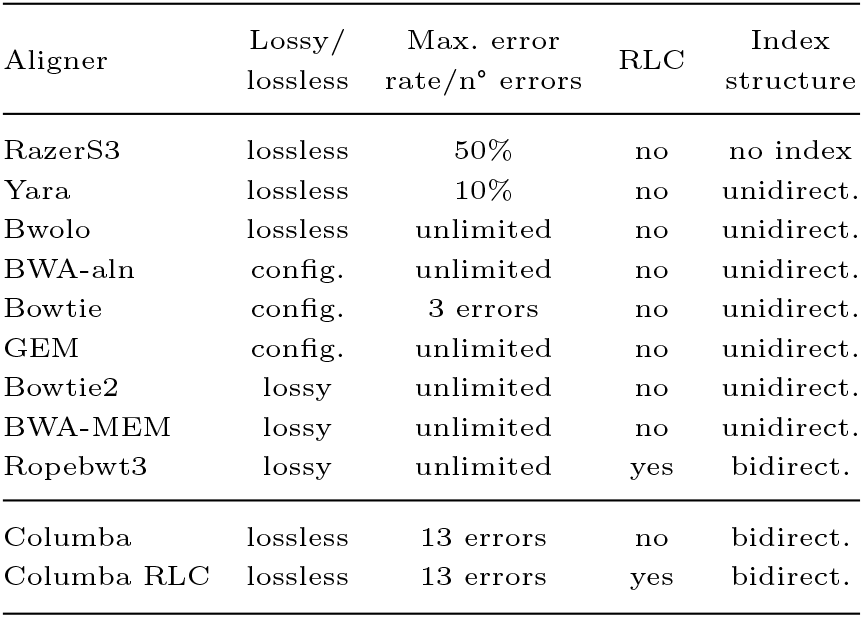
Comparison of aligners. RLC: run-length compression; config.: tool is lossy, but can be configured to perform lossless alignment.

| Aligner | Lossy/<br>lossless | Max. error<br>rate/n° errors | RLC | Index<br>structure |
| --- | --- | --- | --- | --- |
| RazerS3 | lossless | 50% | no | no index |
| Yara | lossless | 10% | no | unidirect. |
| Bwolo | lossless | unlimited | no | unidirect. |
| BWA-aln | config. | unlimited | no | unidirect. |
| Bowtie | config. | 3 errors | no | unidirect. |
| GEM | config. | unlimited | no | unidirect. |
| Bowtie2 | lossy | unlimited | no | unidirect. |
| BWA-MEM | lossy | unlimited | no | unidirect. |
| Ropebwt3 | lossy | unlimited | yes | bidirect. |
| Columba | lossless | 13 errors | no | bidirect. |
| Columba RLC | lossless | 13 errors | yes | bidirect. |

Lossy aligners, such as Bowtie2 (Langmead and Salzberg, 2012) and BWA-MEM (Li, 2013), use a seed-and-extend approach to quickly identify good alignments. However, since seeds –often maximal exact matches, or MEMs– can contain sequencing errors or variants, lossy tools do not guarantee optimal alignments. They typically report only one or a few good alignments per read, rather than providing a comprehensive overview of all possible optimal alignment positions. This limitation can reduce their utility in applications such as bacterial species identification, metagenomics read classification, or Human Leukocyte Antigen (HLA) typing, where reads are aligned against a pan-genome to determine which genome(s) or haplotype(s) best explain the read dataset. In these cases, the pan-genome often contains closely related reference sequences, and an individual read might align with many of them.

In contrast, lossless alignment (also known as *exact* alignment or approximate pattern matching) involves reporting *all* optimal alignments. Optimality is typically defined by the edit (Levenshtein) distance—that is, the total number of substitutions and/or indels in the alignment. Notable examples of lossless aligners include RazerS3 (Weese et al., 2012), Yara (Siragusa, 2015), Bwolo (Vroland et al., 2016), and DREAM-Yara (Dadi et al., 2018), which have their utility in specific applications. For instance, RazerS3 is used in the OptiType tool for HLA typing. Other tools, such as BWA-aln (Li and Durbin, 2009), Bowtie (Langmead et al., 2009), and GEM (Marco-Sola et al., 2012), offer partial support for lossless read alignment, as they often do not report all optimal alignments (see Results). Similarly, Ropebwt3 (Li, 2024), one of the first alignment tools designed for large-scale pan-genome references, reports a single alignment position along with the number of alternative alignment locations, but not their actual positions.

While lossless aligners offer stronger guarantees of optimality, lossy tools are often preferred in practical bioinformatics applications due to their speed. In this paper, we aim to bridge this performance gap by introducing Columba, a high-performance, lossless alignment tool, designed to operate under the edit distance metric. Given single or paired-end reads in FASTQ format and a reference (pan-)genome, Columba reports all optimal alignments in SAM format (Li et al., 2009). It supports two modes:

1. *All mode*: Reports all occurrences of reads within a user-specified edit distance of *k*.
2. *All-best mode*: Reports all optimal of reads occurrences, i.e., occurrences with the smallest edit distance, using a strata-based approach.

Columba can identify alignments with up to *k* = 13 errors per read, making it particularly useful for Illumina sequencing data or for identifying seeds within longer reads. However, it is less suited for spliced aligned of RNA-seq data, as the gap size may exceed the maximum threshold of *k* = 13 errors. For paired-end read input, Columba reports all optimal *proper pairs*. In this context, optimality is defined as the smallest edit distance across both reads. A proper pair is defined by a fragment size that does not deviate by more than six standard deviations from its expected value. The expected fragment size and standard deviation are inferred from the data.

Lossless alignment tools are particularly valuable for pan-genomes, where reads are aligned against a database of many related sequences. Columba can work with two indexes: the bidirectional FM-index and the recently proposed b-move structure (Depuydt et al., 2024). While the FM-index has a memory usage that scales linearly with the total sequence length of the pan-genome, the b-move structure leverages run-length compression (RLC) on the Burrows-Wheeler transform (BWT) to significantly reduce memory requirements. However, this reduction comes at a 30% to 50% cost in performance. This principle of run-length compressing the BWT (Mäkinen and Navarro, 2005) is also the foundation of several recent mapping tools, including MONI (Rossi et al., 2022), PHONI (Boucher et al., 2021), SPUMONI (Ahmed et al., 2021), SPUMONI 2 (Ahmed et al., 2023), and Movi (Zakeri et al., 2024). The key distinction between these tools and Columba is that they focus on MEM-finding, using (pseudo-)matching lengths and statistics, while Columba performs complete alignment.

We demonstrate that Columba (RLC) is significantly faster than existing lossless alignment tools for tasks such as alignment against the human reference genome as well as human and bacterial pan-genomes. Columba is written in C++ (2011 standard), with support for multi-threading. Its source code is available at github.com/biointec/columba under AGPL license.

## Methods

### Search Schemes

Lossless approximate pattern matching (APM) aims to identify all approximate occurrences of a pattern *P* within a text *T* (one or more reference genomes) with up to *k* errors. In this paper, errors can be substitutions, insertions, or deletions, measured using the edit distance metric. By employing a full-text index like the FM-index (Ferragina and Manzini, 2000), a backtracking algorithm systematically explores all candidate occurrences character by character. However, the search tree induced by this method quickly becomes impractically large, even for moderate values of *k*.

Columba employs *search schemes* (Kucherov et al., 2014) and a bidirectional full-text index to significantly reduce the size of the search tree. The pattern *P* is divided into *p* potentially uneven parts *P*_*i*_ (*i* = 1, …, *p*). A search scheme 𝒮 consists of multiple searches *S*, where each search is represented by the triplet *S* = (*π, L, U*), where:

- *π* is a permutation of {1, 2, …, *p*} that specifies the order in which the parts of *P* are matched using the index, i.e., *P*_*π*[1]_, *P*_*π*[2]_, …, *P*_*π*[*p*]_. It must satisfy the *connectivity property*, ensuring that partial matches are extended contiguously, either to the left or to the right.
- *L* and *U* are arrays of size *p*, where *L*[*i*] and *U* [*i*] respectively denote the lower and upper bounds on the cumulative number of errors after part *P*_*π*[*i*]_ has been processed. The values of *L* and *U* must increase monotonically and *L*[*i*] ≤ *U* [*i*], for all *i*.

The searches in a search scheme are designed to collectively cover all possible distributions of *k* errors across the parts of *P*. Therefore, each approximate occurrence of *P* is guaranteed to be identified by at least one search. The lower and upper bounds defined by the *L* and *U* arrays help reduce the search space and limit the redundant reporting of occurrences across different searches. Such redundant occurrences are filtered in a post-processing step.

Designing efficient search schemes is non-trivial, especially for higher values of *k*. The pigeonhole principle forms the basis for some of the simplest search schemes (Lam et al., 2009), where each search targets one part of *P* for exact matching, and allows up to *k* errors in the remaining parts. More efficient schemes, such as those proposed by Kucherov et al. (2014), Kianfar et al. (2017), and Pockrandt (2019), introduce tighter error bounds to make the overall process more efficient. In this work, we use the MinU search schemes recently proposed by us (Renders et al., 2024). The MinU schemes were designed using an Integer Linear Programming (ILP) approach for up to *k* = 7 errors and a greedy algorithm for *k* = 8 … 13 errors. The MinU schemes were demonstrated to be very efficient for practical applications.

### Bidirectional FM-Index and move structure

The FM-index supports only backward search: given a partial match *M*, occurrences of *cM* (where *c* is a character) can be identified in *O*(1) time (Ferragina and Manzini, 2000). However, search schemes require *bidirectional* search functionality, as this allows for matching some middle part of *P*, followed by left and right extensions in an arbitrary order. The bidirectional FM-index, introduced by Lam et al. (2009), allows for extending a partial match *M* to either *cM* or *Mc* in *O*(*σ*) time, with *σ* the size of the alphabet (*σ* = 4 for DNA sequences). Columba implements bidirectional character extensions in *O*(1) time using a fast algorithm by Pockrandt et al. (2017).

For large pan-genomes, the bidirectional FM-index can have prohibitively large memory requirements. In such cases, the BWT, a key component of the FM-index, often contains long runs –consecutive occurrences of the same character– making it inherently compressible. To support lossless approximate pattern matching in run-length compressed space, Columba RLC replaces the bidirectional FM-index with the bidirectional *move structure* (b-move in short) (Depuydt et al., 2024). The *r*-index was the first index to support both counting and locating exact occurrences of patterns in *O*(*r*) space, where *r* is the number of runs in the BWT (Gagie et al., 2018, 2020). The move structure, proposed by Nishimoto and Tabei (2021), forms an alternative to the *r*-index with significantly better performance in practice. b-move extends the move structure to enable bidirectional character extensions in a conceptually similar manner to the bidirectional FM-index. Like the move structure, b-move supports fast, cache-efficient character extensions in run-length compressed space.

To construct the suffix array, Columba employs either `libsais`^1^ or `libdivsufsort`^2^, depending on the length of the text. In the Columba RLC, there are two methods for index construction: an in-memory approach; which is fast but requires substantial RAM, and a prefix-free parsing method (Boucher et al., 2018), which is slower but significantly more memory-efficient.

### Performance enhancements

Columba incorporates several performance enhancements. Recall that search schemes partition search patterns into *p* parts. Instead of using static partitioning (e.g., equally-sized parts), we developed a *dynamic partitioning* method. This approach tailors the partition sizes for each pattern individually, ensuring a balanced computational workload across searches within the search scheme. The workload for each search is estimated by the number of exact matches of the part *P*_*i*_ that is matched first by that search. For example, if part *P*_*i*_ has a large number of exact matches in *T*, any search continuing from that part would likely generate an excessively large search tree to explore. By increasing the size *P*_*i*_, its number of exact matches decreases, thereby reducing its search tree and associated runtime. Dynamic partitioning of search patterns reduces runtime by roughly 25% (Renders et al., 2021).

To further optimize performance, we implemented a *dynamic selection* mechanism for search schemes. For a given number of errors *k*, multiple search schemes may be available, and their efficiency can vary depending on the specific search pattern. The dynamic selection mechanism uses a heuristic to identify the search scheme with the lowest estimated workload, again relying on the number of exact matches of each part *P*_*i*_. This approach further reduces runtime by approximately 10% (Renders et al., 2024).

The use of the edit distance metric inherently involves some redundancy: a pattern *P* may be reported multiple times with slightly different start and/or end positions in *T*. For example, a leading or trailing gap can be exchanged for a substitution. This redundancy becomes particularly problematic in the context of search schemes, where parts are matched individually, and direction switches may occur. To address this issue and avoid redundant calculation, we applied a clustering technique to focus computations on representative alignment paths while maintaining the lossless nature of APM (Renders et al., 2021). Additionally, we implemented a bit-parallel approach, inspired by Myers (1998) and Hyyrö (2003), to compute all edit distance values in a row of a banded alignment matrix in simultaneously.

Finally, while Columba primarily relies on a full-text index for APM, it dynamically switches to in-text verification once the number of candidate occurrences drops below a threshold (4 by default). In-text verification is performed in a bit-parallel, cache-efficient manner, avoiding the need for more costly character-by-character extensions using the index (Renders et al., 2022).

## Material and Experimental Setup

The benchmarks on a single human genome, the bacterial pan-genome and for HLA-typing were obtained using a single thread of a 64-core Intel Xeon E5-2698 v3 CPU, operating at a base clock speed of 2.30 GHz, with 256 GiB of available RAM. For the benchmark on the human pan-genome, a single thread of a 64-core AMD EPYC 7773X (Milan-X @ 2.2 GHz) processor was used, with 940 GiB of RAM available. All experiments were run 10 times; the standard deviation was always lower than 1.7%. The lossless tools use the edit distance.

### Comparing the output of different aligners

The aligners report results in SAM format (Li et al., 2009), except for Ropebwt3, which reports in PAF format^3^. Ropebwt3 only reports 1 alignment per read along with the number of alignments BWA-aln outputs alignments in the intermediate SAI format and requires BWA-SAMSE to convert its output to SAM format.

Comparing the output of various aligners is challenging, even when using the SAM format. We examine all equally best alignments per read, however not all aligners restrict their output to the best alignment; for example, BWA and Bowtie sometimes report additional alignments with a worse score. Additionally, supplementary co-optimal alignments are handled differently: Columba lists them as separate records, while BWA uses the XA tag. Yara also uses the XA tag but in a different format, though it can output separate records for supplementary alignments without CIGAR strings, which complicates SAMtools compatibility. Similarly, Columba RLC never provides a CIGAR string, while Columba always reports a CIGAR string.

The interpretation of maximum error rate (or minimum identity score) can slightly vary across the different tools, resulting in tiny variations in the output. Finally, slight variations in alignment positions under an edit distance model necessitate some manual inspection, as aligners may produce subtly different results for the same region.

## Results

Alignment of Illumina reads to the human reference genome We aligned 1 million Illumina NovaSeq 6000 reads (151 bp), randomly sampled from a whole genome sequencing dataset (accession no. SRR9091899), to the hg38 human reference genome (Schneider et al., 2017). In this initial benchmark, the reads were treated as unpaired. We used both lossless and lossy alignment tools. For lossless alignment, we included Columba, Yara, BWA-aln with the -N flag, and Bowtie with the -a flag; these flags enable lossless search. For lossy alignment, we used BWA-MEM, Bowtie2, and Bowtie2 in ‘very sensitive’ (VS) mode. The lossless alignment tools were configured to report all optimal alignments in *all-best* mode for maximum error rates ranging from 0-8%. The command line options for each tool are listed in Suppl. Data Section 1. Table 2 shows the runtime, peak memory usage, and the percentage of reads that aligned by each tool.

**Table 2.** Runtime and peak memory usage of lossless alignment tools (Columba, Yara, BWA in aln mode, and Bowtie) at varying maximum error rate thresholds, and lossy alignment tools (BWA-MEM and Bowtie2 in normal and very sensitive (VS) modes), for aligning 1 000 000 Illumina reads (length 151 bp) from a larger WGS dataset to the human reference genome using a single thread. The alignment percentage of reads is noted alongside each runtime.

| Tool | Peak<br>Memory | Maximum Error Rate |  |  |  |  |
| --- | --- | --- | --- | --- | --- | --- |
|  |  | 0% | 2% | 4% | 6% | 8% |
| Lossless alignment tools |  |  |  |  |  |  |
| Columba (this paper) | 11.06 GB | 24s (66.6%) | 45s (91.1%) | 1m 27s (95.2%) | 4m 33s (96.8%) | 21m 09s (97.7%) |
| Yara | 4.94 GB | 33s (66.6%) | 1m 24s (91.1%) | 11m 42s (95.2%) | 1h 57m 10s (96.8%) | 11h 11m 42s (97.7%) |
| BWA-aln | 4.51 GB | 1m 2s (66.6%) | 41m 16s (91.1%) | 3h 38m 9s (95.0%) | 8h 41m 28s (96.5%) | 13h 55m 54s (97.2%) |
| Bowtie | 2.29 GB | 37s (66.6%) | 2m 28s (89.9%) | not supported | not supported | not supported |
| Lossy alignment tools |  |  |  | All Error Rates |  |  |
| BWA-MEM | 5.19 GB | 6m 46s (99.8%) |  |  |  |  |
| Bowtie2 | 3.22 GB | 7m 53s (98.8%) |  |  |  |  |
| Bowtie2 (VS) | 3.23 GB | 17m 01s (99.0%) |  |  |  |  |

In terms of performance, Columba significantly outperforms existing lossless aligners. It is 1.4×, 1.9×, 8.1×, 25.8×, and 37.8× faster than Yara, its closest competitor, at error rates of 0%, 2%, 4%, 6%, and 8%, respectively. This indicates that optimized search schemes offer superior performance for lossless approximate pattern matching, particularly at higher error rates. The runtimes also indicate that BWA-aln was not primarily designed for lossless alignment, consistent with its manual, which notes that the -N flag may lead to significantly lower performance. Bowtie with the -a flag is slower than Yara and Columba at error rates 0 and 2%, and it does not support lossless alignment beyond 3 errors, which corresponds to a 2% error rate for this dataset.

When comparing Columba to lossy tools, we observe that up to a maximum error rate of 6% (a reasonable threshold for Illumina data), Columba is faster than the lossy alternatives. Even at an error rate of 8%, Columba’s runtime remains competitive with Bowtie2 in VS mode. Due to its use of a bidirectional index, Columba requires more RAM than other tools. However, at 11.06 GB, it still fits comfortably on a modern laptop.

In principle, all lossless alignment tools should produce optimal –and therefore identical– results. Columba and Yara produce near-identical results, with minor differences arising from a slightly different interpretation of the error rate, handling of ‘N’ characters and treatment of overlapping alignments. For example, one tool may list overlapping alignments as separate alignments, while others may flag one as redundant. In contrast, despite being configured as lossless tools, BWA-aln and Bowtie do not always report the optimal alignment(s), as evidenced by slightly lower alignment percentages (Table 2). Additionally, when multiple optimal alignments exists for a read, often only one or a few are reported.

On the other hand, the lossy alignment tools align a higher fraction of reads. For example, at a 6% error rate threshold, Columba mapped 96.8% of the reads, while BWA-MEM found alignments for nearly all (99.8%) reads. Upon inspection, these additional alignments contain error rates *>* 6%, often due to spliced alignment or read clipping.

In Suppl. Data Section 2, we repeated the same benchmark, this time aligning 1 million reads pairs (2 million reads in total). This task is significantly more challenging because optimal proper pairs are not necessarily obtained by combining optimal alignments of the individual reads. Nevertheless, Columba outperforms all lossless aligners by a large margin, and is competitive with lossy aligners until an error rate of 6%.

### HLA Typing from NGS data using OptiType

Lossless alignment tools are crucial for certain bioinformatics applications, including Human Leukocyte Antigen (HLA) typing from next-generation sequencing (NGS) data. Accurate, high-resolution HLA typing is essential for understanding immune system function and has significant applications in organ transplantation compatibility, autoimmune disease research, and cancer immunotherapy.

In a recent survey paper on HLA typing tools (Claeys et al., 2023), OptiType (Szolek et al., 2014) emerged as one of the most accurate tools for predicting HLA genotypes from NGS data. It focuses on Major Histocompatibility Complex (MHC) Class I alleles, including HLA-A, HLA-B, and HLA-C. For each read, OptiType requires a complete list of all possible candidate alleles from which the read could have originated. To achieve this, OptiType uses the lossless aligner RazerS3 (Weese et al., 2009, 2012) to map reads to a comprehensive collection of HLA allele sequences. Next, leveraging an integer linear programming (ILP) approach, OptiType identifies the most likely set of HLA alleles that explain the sequencing data.

While OptiType delivers precise genotyping, it was also found to be computationally intensive (Claeys et al., 2023), partly due to the alignment step. To address this, we use Columba as a drop-in replacement for RazerS3, which required only minimal changes to OptiType’s script (see Suppl. Data Section 3). Using the dataset from the survey paper, we evaluated OptiType by aligning slices of 1 012 CRAM files of whole-exome sequencing (WES) data from the 1000 Genomes on GRCh38 project (Zheng-Bradley et al., 2017), to an HLA-gene reference database^4^ (Szolek et al., 2014).

Fig. 1 shows the runtime distribution of the alignment phase for both Columba and RazerS3 across the 1 012 samples. In both cases, 16 CPU cores were used. Columba achieved a median runtime of 4.83 s compared to 57.96 s for RazerS3, yielding a 12× speedup. Consequently, using Columba reduced the overall runtime of the OptiType pipeline (including the ILP step) by 44% (median value). As Columba and RazerS3 produce near-identical output, OptiType’s accuracy remains unaffected. We further evaluated OptiType on 373 RNA-seq samples (see Suppl. Data Section 3). In this case, the median runtime of the alignment phase is decreased from 4.82 minutes (RazerS3) to 58 seconds (Columba), demonstrating significant efficiency gains for RNA-seq data as well.

**Fig. 1:**
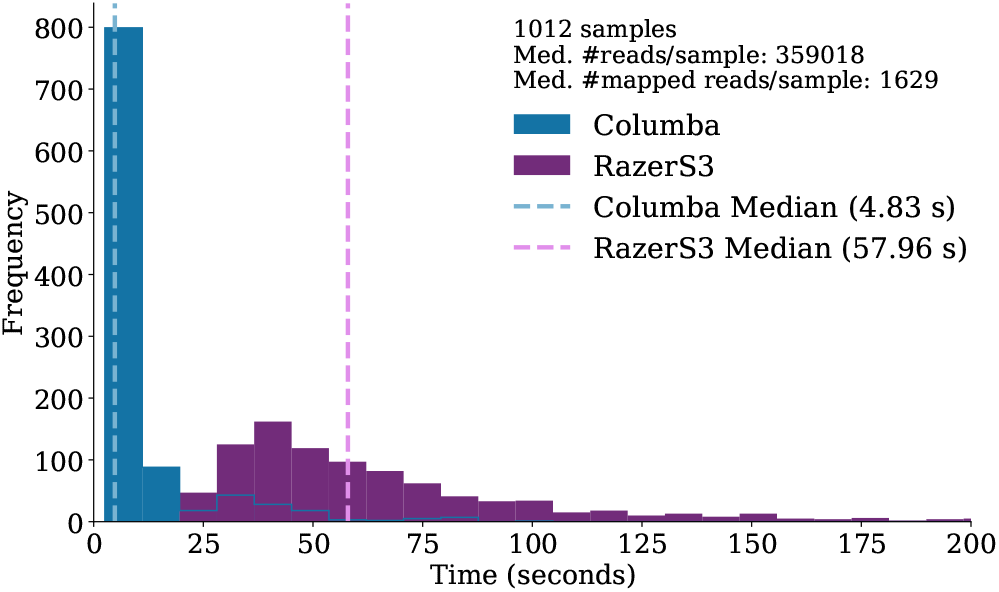
Runtime distribution of OptiType’s alignment phase using Columba and RazerS3 accross 1 012 samples. The x-axis is capped at 200 seconds. RazerS3’s distribution continues beyond this value.

### Alignment to bacterial pan-genomes

We aligned 1 million *Escherichia coli* reads (150 bp, Illumina NovaSeq 6000), randomly sampled from a larger sequencing dataset (SRR19479103), to a pan-genome comprising 6 115 bacterial strains: 2 594 *Escherichia coli*, 1 183 *Salmonella enterica*, 944 *Staphylococcus aureus*, 637 *Pseudomonas aeruginosa*, 254 *Bacillus subtilis*, 255 *Listeria monocytogenes*, 203 *Enterococcus faecalis*, and 45 *Limosilactobacillus fermentum* genomes (∼28 billion bp in total, accessions numbers in Suppl. Data Section 4) to evaluate runtime and peak memory usage. The lossless alignment tools were again run in *all-best* mode, a configuration suited for pan-genome analysis as it provides a comprehensive overview of all possible genomes a read could have originated from. This is particularly relevant for tasks such as bacterial species or strain identification and taxonomic sequence classification for metagenomics. The comparison includes Columba RLC, the run-length compressed variant of Columba that employs the bidirectional move structure instead of the bidirectional FM-index, as well as Ropebwt3, a mapper specifically designed for pan-genome references.

Table 3 shows that Columba is the fastest tool among the evaluated lossless aligners, outperforming Yara by up to 10.1×. Columba and Columba RLC generate identical alignments as they share the same alignment methodology, differing only in index structure. Because Columba RLC does not store the reference genome in memory, its SAM output does not contain the CIGAR string. The outputs of Yara and Columba (RLC) are again nearly identical, with minor discrepancies caused by small implementation differences. As before, BWA-aln, despite the use of the -N flag, does not always report an optimal alignment for each read. This is reflected in the slightly lower alignment percentage. Additionally, BWA-aln does not comprehensively report all optimal alignments, excluding nearly 312 million co-optimal alignments, over 99.7% of those reported by Columba and Yara. In the context of this pan-genome, many reads have hundreds of optimal alignments, and BWA-aln typically only one or a few of those. This vastly reduced output volume explains BWA-aln’s lower runtime on this dataset at a 0% error rate.

**Table 3.** Runtime and peak memory usage of lossless alignment tools (Columba, Columba RLC, Yara, and BWA in aln mode) at varying maximum error rate thresholds, and lossy alignment tools (BWA-MEM and Ropebwt3), for aligning 1 million *E. coli* Illumina reads (length 150 bp) to a pan-genome comprising 6 115 bacterial strains using a single thread. The alignment percentage of reads is noted alongside each runtime.

| Tool | Peak<br>Memory | Maximum Error Rate |  |  |  |  |
| --- | --- | --- | --- | --- | --- | --- |
|  |  | 0% | 2% | 4% | 6% | 8% |
| Lossless alignment tools |  |  |  |  |  |  |
| Columba | 86.00 GB | 10m 14s (75.3%) | 12m 9s (94.6%) | 12m 26s (96.6%) | 13m 46s (97.4%) | 18m 12s (97.8%) |
| Columba RLC | 14.21 GB | 15m 16s (75.3%) | 18m 39s (94.6%) | 19m 31s (96.6%) | 21m 10s (97.4%) | 25m 23s (97.8%) |
| Yara | 48.27 GB | 11m 55s (75.3%) | 19m 10s (94.6%) | 26m 24s (96.6%) | 38m 54s (97.4%) | 3h 4m 43s (97.8%) |
| BWA-aln | 39.35 GB | 3m 60s (75.3%) | 37m 39s (94.5%) | 1h 11m 41s (96.5%) | 2h 30m 35s (97.2%) | 4h 35m 1s (97.6%) |
| Lossy tools |  |  |  | All Error Rates |  |  |
| BWA-MEM | 46.92 GB | 30m 13s (99.2%) |  |  |  |  |
| Ropebwt3 | 2.67 GB | 42m 9s (99.2%) |  |  |  |  |

Compared to Columba, Columba RLC reduces memory usage by 5.6× (see Table 3). Note that Columba RLC improves the memory complexity of Columba from O(*n*) (with *n* being the total size of the reference (pan-)genome) to O(*r*) (with *r* being the number of runs in the BWT). Therefore, the relative memory gain will only increase for larger (and more repetitive) pan-genomes. However, this memory efficiency comes at the cost of a 1.4× to 1.6× runtime increase compared to Columba. Even so, Columba RLC remains faster than other lossless and lossy alignment tools. Ropebwt3 achieves superior compression compared to Columba RLC but at the expense of significantly longer runtimes. Columba RLC is nearly two times faster for an error rate of 6% and 40% faster for an error rate of 8%. Importantly, Ropebwt3 reports only a single alignment per read along with the count of equally optimal hits (without reporting their positions). For 95.9% of the reads, the alignment produced by Ropebwt3 matched one of the alignments reported by Columba (for an error rate 8%) within at most 50 bp. For 1.8% of the reads, Ropebwt3 reported alignments beyond an error rate of 8% (12 errors), which Columba did not produce. However, for 9 526 reads (0.95%), Columba reported more alignments (3 800 953 alignments not reported by Ropebwt3), while Ropebwt3 produced a higher number of alignments for 4 527 reads (resulting in 279 042 additional alignments). As Columba and Yara are lossless and produce near-identical output, we infer that the additional alignments reported by Ropebwt3 correspond to overlapping occurrences in the reference genome (less than 20 bp apart), which are considered spurious by Columba and Yara.

Supplementary Fig. 1 illustrates the runtimes for Columba (RLC), Yara, Bowtie2, BWA-MEM, and Ropebwt3 when utilizing multiple threads. The results demonstrate that all tools benefit from multi-threading. However, at higher thread counts, the performance of the lossless tools starts to be constrained by the speed at which output can be written to disk.

### Alignment to Human Pan-genome Reference

We aligned the same 1 million Illumina reads from the first benchmark, but now against pan-genomes containing up to 64 human haplotypes, sourced from the 1-year freeze of the Human Pangenome Consortium, which provides a comprehensive representation of genetic diversity (Liao et al., 2023). Indexes were constructed for Columba, Columba RLC, and Ropebwt3. We could evaluate Columba for only up to 16 human haplotypes, as constructing the index for 32 haplotypes exceeded the available RAM on the machine (940 GiB). Columba and Columba RLC were executed in *all-best* mode with a maximum error rate of 6%, while Ropebwt3 was used with its default settings.

Fig. 2 shows that Columba’s memory usage scales linearly with the number of haplotypes, as shown by the trendline. In contrast, Columba RLC and Ropebwt3 demonstrate more favorable memory requirements. Even though both leverage run-length compression on the BWT and share the same *O*(*r*) memory complexity, Columba RLC’s bidirectional move structure implementation introduces a larger constant prefactor. As a result, Ropebwt3 has a better memory footprint than Columba RLC in practice. In comparison to Columba, Columba RLC exhibits lower memory usage only when processing more than 16 haplotypes.

**Fig. 2:**
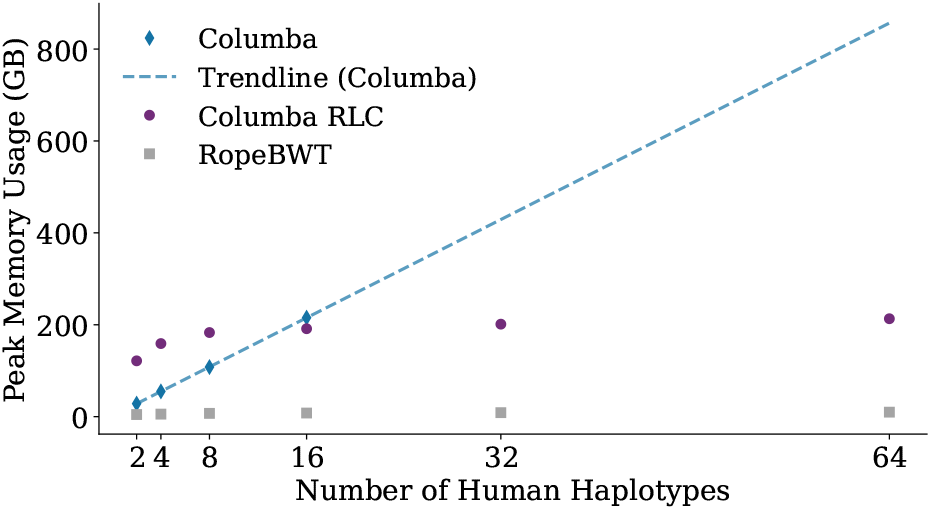
Comparison of peak memory usage among Columba, Columba RLC, and Ropebwt3 for aligning 1 000 000 reads to a pan-genome, with varying number of human haplotypes.

Fig. 3 illustrates the runtime for aligning 1 million reads to the human pan-genomes. Columba RLC has a performance penalty compared to Columba but remains significantly faster than Ropebwt3. The increase in runtime for Columba (RLC) with larger pan-genome sizes can be attributed to the greater volume of output that needs to be produced, i.e., a larger number of optimal alignments. In contrast, Ropebwt3 reports only a single alignment and the number of optimal alignments.

**Fig. 3:**
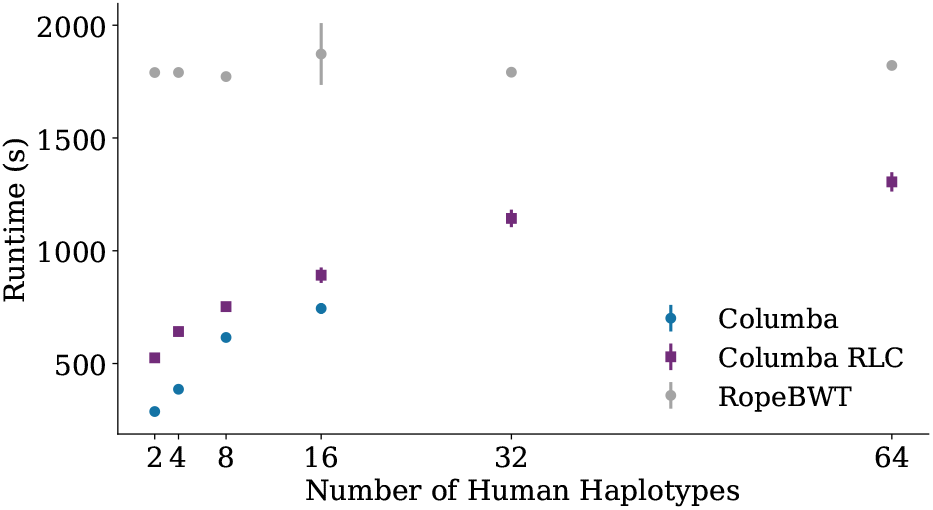
Comparison of runtime among Columba, Columba RLC, and Ropebwt3 for aligning 1M reads to a pan-genome, with varying number of human haplotypes.

When comparing the output of Columba (RLC) and Ropebtw3, a similar analysis can be made as for the bacterial pan-genome.

For 956 398 reads (95.6%), both tools report the same number of alignments, with each alignment identified by Ropebwt3 also present in Columba’s output. For 20 187 reads (2.0%), Ropebwt3 provides an alignment while Columba does not, because the error rate exceeds the 6% threshold. On the other hand, for 16 reads Columba reports alignment(s), while Ropebwt3 does not report any alignments For 10 474 reads (1.0%), Ropebwt3 did not report (any of) the alignment(s) with the lowest edit distance, while for 6 871 reads, Columba reported additional alignments, leading to a total of 7 601 497 alignment counts not reported by Ropebwt3. Conversely, Ropebwt3 reported a higher number of alignments for 4 371 reads then Columba did. Since Ropebwt3 does not report detailed alignments but rather provides aggregate counts, it is challenging to ascertain why these alignments are not corroborated by Columba. It is plausible that Ropebwt3 either produces spurious alignment counts (e.g., alignments that are overlapping) or includes misreported results.

## Discussion and Conclusion

We propose Columba, a fast and feature-rich lossless alignment tool. It accepts FASTQ or FASTA files as input and produces SAM records as output, with support for multi-threading to enhance performance.

Unlike widely used lossy aligners such as BWA-MEM and Bowtie, Columba guarantees optimality and completeness of its output. Optimality is evaluated in terms of edit distance, a standard metric for assessing differences between reads and reference sequences. However, certain applications may benefit from alternative scoring schemes, such as affine gap penalties, which are likely more biologically relevant. In *all* mode, completeness means that all occurrences in the reference sequences with up to a predefined number of errors (*k*) are reported, with support for up to *k* = 13 errors. In *all-best* mode, completeness refers to reporting all optimal occurrences (i.e., occurrences with the lowest edit distance to the query read). Columba supports paired-end read mapping; in this mode, only alignment positions that conform to the expected fragment size are reported. If no such proper pair can be found, the reads are treated as unpaired. Our evaluation demonstrates that Columba is significantly faster than other lossless alignment tools such as RazerS3 and Yara, especially when aligning reads with a higher number of errors. This performance improvement is achieved through optimized search schemes and an efficient bit-parallel implementation. However, as the search schemes require a bidirectional index, Columba’s memory requirements are generally double those of tools that use a unidirectional index.

Columba supports both single-genome and pan-genome references. For pan-genomes, Columba RLC leverages the bidirectional move structure, a run-length compressed index. This approach significantly reduces memory usage when multiple similar genomes are included in the pan-genome, albeit with a slight trade-off in speed. Other tools such as Ropebwt3 offer a different time-space tradeoff: Ropebwt3 is able to better compress pan-genomes, but appears slower. Additionally, Ropebwt3 does not report all alignments, only the alignment count.

Columba is a robust and versatile alignment tool for a wide range of applications. Its successful integration into the OptiType pipeline for HLA genotyping highlights its practical utility and effectiveness. We believe Columba and Columba RLC can also be relevant for applications such as metagenomics read classification and bacterial species or strain identification; or pan-genome graph alignment and subsequent visualization (Depuydt et al., 2023). Finally, Columba (RLC) can be used to benchmark the output of lossy alignment tools.

## Author contributions statement

The authors wish it to be known that, in their opinion, the first two authors should be regarded as joint first authors. **Luca Renders**: Conceptualization, Methodology, Software, Validation, Investigation, Writing - Original Draft. **Lore Depuydt**: Software, Methodology, Validation, Writing - Review & Editing. **Travis Gagie**: Conceptualization, Writing - Review & Editing. **Jan Fostier**: Conceptualization, Supervision, Writing - Review & Editing.

## Acknowledgments

This work was supported by the Research Foundation – Flanders (FWO) through a PhD Fellowship SB [grant number 1SE7822N] to L.R. and a PhD Fellowship FR [grant number 1117322N] to L.D.

## Footnotes

1 https://github.com/IlyaGrebnov/libsais

2 https://github.com/y-256/libdivsufsort

3 https://github.com/lh3/miniasm/blob/master/PAF.md

4 https://raw.githubusercontent.com/FRED-2/OptiType/refs/heads/master/data/hla_reference_dna.fasta

