## Supplementary Material for "Columba: Fast Approximate Pattern Matching with Optimized Search Schemes"

#### 1 Commands Used for the Different Tools

Table 1: Commands (except I/O) used in the alignment benchmarks. $E$ stands for the maximal error rate and $L$ stands for the average length of the reads (151 bp for the benchmarks on the human reference genome and 250 bp for the benchmarks on the *listeria monocytogenes* pan-genome.

| Tool | Command |
|---|---|
| Yara | `./yara_mapper -y full -t 1 -e [E] -sa record` |
| RazerS3 | `./razers3 -i [100 - E] -m 99999 -dr 0` |
| BWA-aln | `./bwa aln -N -n [` $1 - \sum_{i=0}^{\lfloor \frac{L \cdot E}{100} \rfloor - 1} \frac{e^{-\lambda} \cdot \lambda^i}{i!}$ `]` with, $e = 0.02$, or `./bwa aln -N -n 0` if $E = 0$, followed by `./bwa samse` |
| Bowtie | `./bowtie -a --best --strata -v [E·L]` |
| Columba | `./columba -I [100-E]` |
| BWA-MEM | `./bwa mem -L 10000` |
| Bowtie2 | `./bowtie2` |
| Bowtie2 (Very Sensitive) | `./bowtie2 --very-sensitive` |
| Ropebwt3 | `./ropebwt3 sw -t1` |

### 2 Timings for Paired-end alignment to HRG

Table 2: Runtime and peak memory usage of lossless alignment tools (Columba, Yara, BWA in aln mode, and Bowtie) at varying maximum error rate thresholds, and lossy alignment tools (BWA-MEM and Bowtie2 in normal and very sensitive (VS) modes), for aligning 1 000 000 pairs of Illumina reads (length 151 bp) from a larger WGS dataset to the human reference genome using a single thread. The alignment percentage of paired reads (reads mapped in proper pair) is noted alongside each runtime.

| **Tool** | **Maximum Error Rate** | | | | |
|---|---|---|---|---|---|
| | 0% | 2% | 4% | 6% | 8% |
| **Lossless alignment tools** | | | | | |
| **Columba** | **44s** (45.4%) | **1m 57s** (82.7%) | **5m 01s** (89.8%) | **19m 54s** (92.9%) | **1h 49m 21s** (94.7%) |
| Yara | 11m 04s (42.0%) | 7m 14s (79.1%) | 29m 36s (85.5%) | 4h 32m 07s (88.2%) | 1d 00h 55m 19s (89.8%) |
| BWA-aln | 2m 42s (73.0%) | 1h 22m 42s (89.8%) | 7h 28m 21s (94.5%) | 17h 01m 55s (94.6%) | 1d 02h 50m 36s (94.6%) |
| Bowtie | **46s** (7.4%) | 1h 11m 10s (11.2%) | not supported | not supported | not supported |
| **Lossy alignment tools** | | | **One Timing** | | |
| BWA-MEM | | | 16m 59s (99.5%) | | |
| Bowtie2 | | | 13m 55s (84.5%) | | |
| Bowtie2 (VS) | | | 25m 45s (84.7%) | | |

Among lossless aligners, Columba consistently proves to be the fastest, completing the alignment in just 19m 54s at a 6% error rate, significantly outperforming Yara (4h 32m 07s) and BWA-aln (17h 01m 55s). Moreover, Columba's runtime remains competitive with lossy alternatives. The comparison of output across aligners is complicated by the distinct methods each tool uses to define "proper pair" mappings. Columba dynamically infers the mean fragment size and orientation, reporting pairs within six standard deviations of the detected mean fragment size. When multiple such pairs are possible, it selects those with the minimal combined edit distance.

The significantly higher alignment percentage observed for BWA-aln lower error rates can be attributed to its inclusion of alignments with clippings and its occasional allowance of errors in the mate, even when the maximum edit distance is strictly set to 0. Conversely, the significantly lower alignment percentage observed for Bowtie (both versions 1 and 2) can be attributed to its use of a fixed insert size, with default values of 250 and 500, respectively.

### 3 HLA Typing Experiment

#### 3.1 Runtime Histogram for RNA Typing

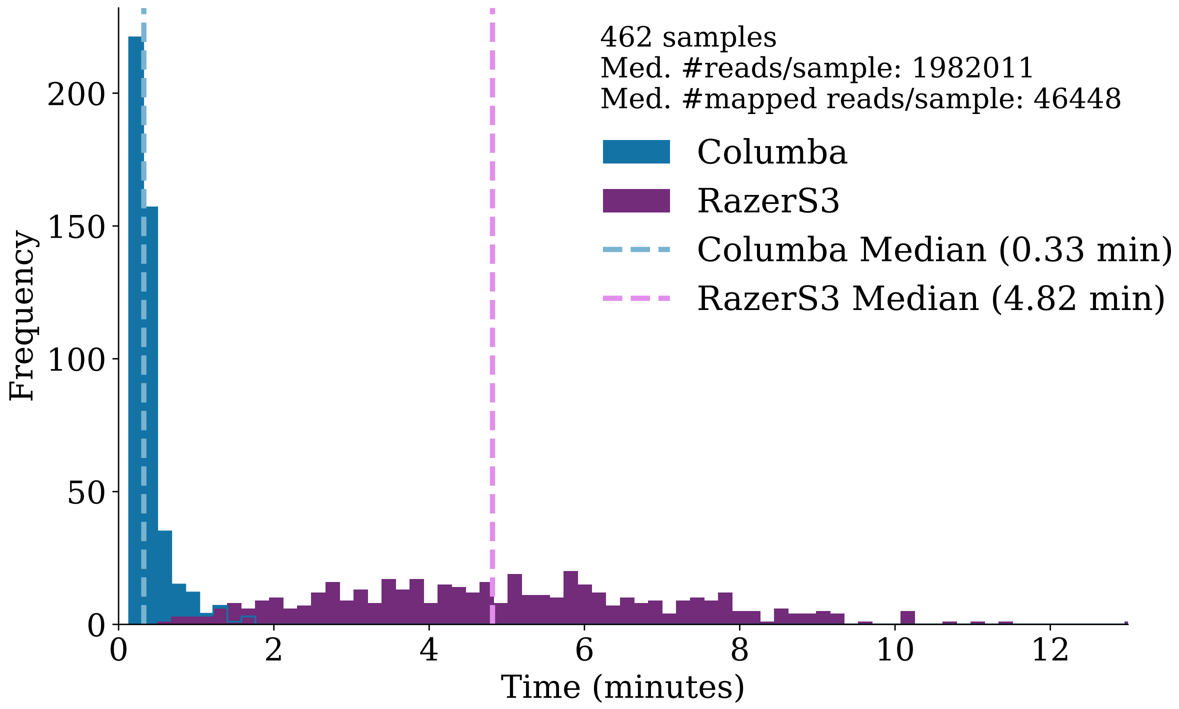


Figure 1: Comparison of the distribution of runtimes of the alignment phase of the Optitype pipeline with Columba and RazerS3 as the aligner for RNA sequences. Note that the figure is zoomed in on the range of 0 to 13 minutes, and that the distribution of RazerS3 has a tail that continues beyond this range.

#### 3.2 Drop-in Nature of Columba in Optitype Script

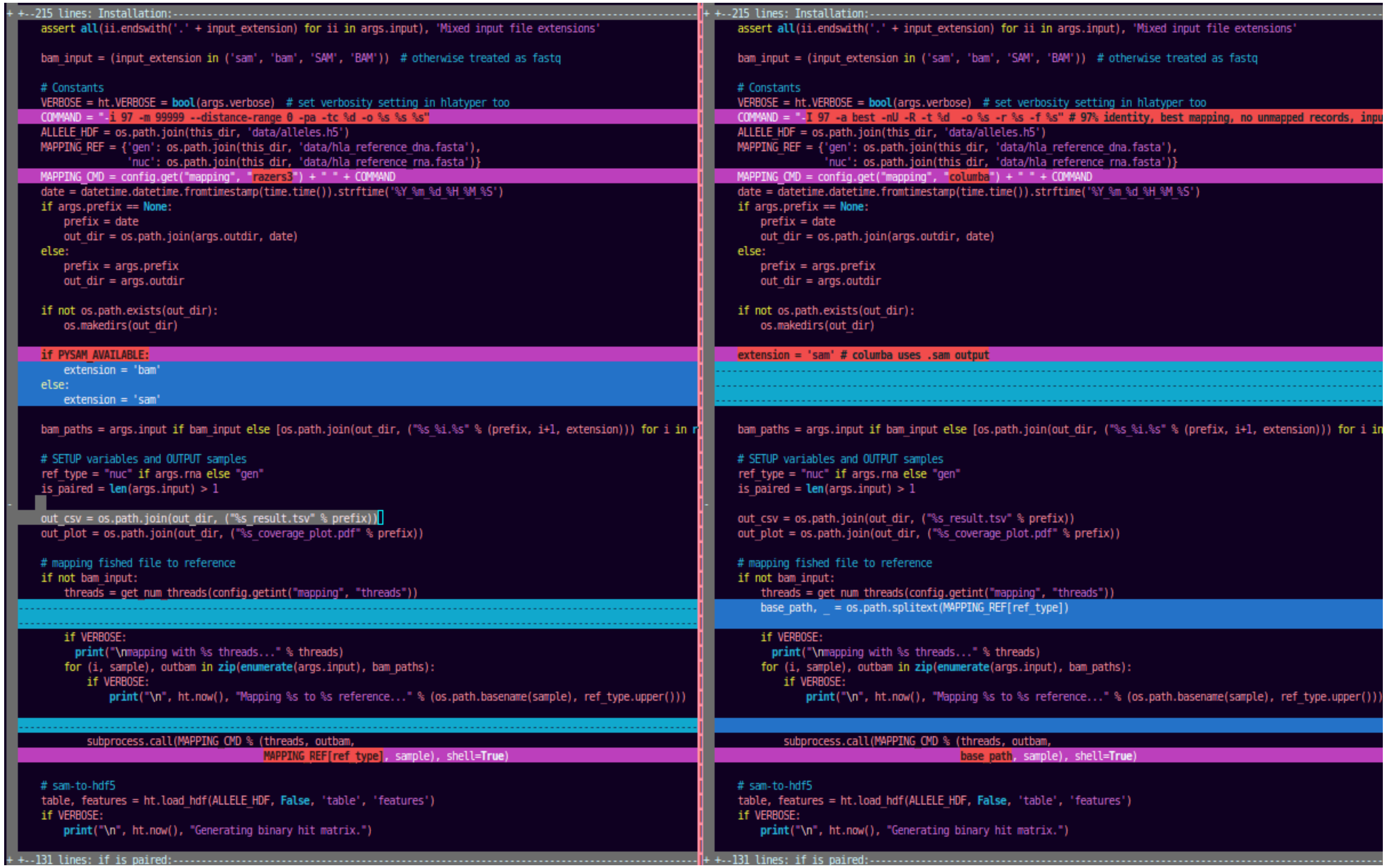

Figure 2: Illustration of Columba's drop-in compatibility in the Optitype script. The figure shows a side-by-side comparison of the two Optitype script versions illustrating the drop-in nature of replacing the razers3 mapping tool with columba for HLA typing. The left panel shows the original script utilizing Razers3 with specific command-line parameters for mapping, while the right panel demonstrates the modified version. This visual highlights the minimal modifications required to adapt existing workflows for Columba, showcasing its compatibility and flexibility in the pipeline.

### 4 Accession numbers for the bacterial pan-genome dataset

The accession numbers for the 6115 bacterial genomes are provided in the supplementary file `BacterialPanGenomeData.tsv`.

### 5 Multi-threaded Timings for the Listeria Monocytogenes pan-genome

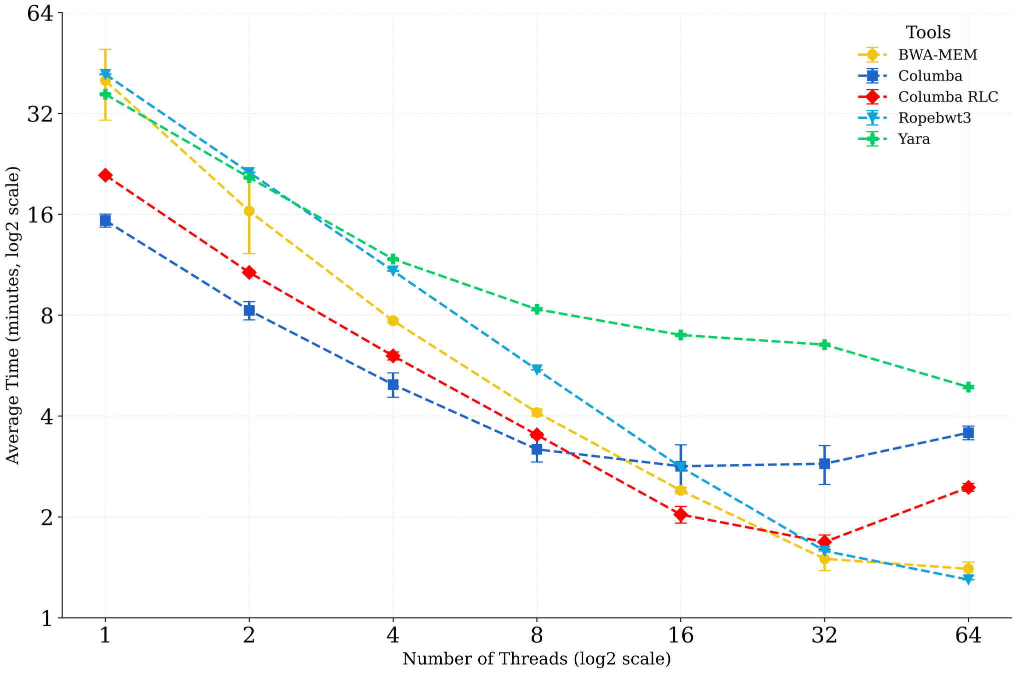


Figure 3: Multi-threaded timings for the benchmark where 1 reads are aligned to the pangenome consisting of 6115 bacterial genomes with various tools. The lossless tools (Yara and Columba) are configured with an allowed error rate of 5%. The lossless tools report significantly more occurrences than lossy aligners, leading to contention and throttling at higher thread counts due to disk write speed becoming a bottleneck.
